# A *Recql5* mutant enables complex chromosomal engineering of mouse zygotes

**DOI:** 10.1101/2023.04.06.535871

**Authors:** Satoru Iwata, Miki Nagahara, Takashi Iwamoto

**Affiliations:** Center for Education in Laboratory Animal Research, Chubu University; Department of Biomedical Sciences, College of Life and Health Sciences, Chubu University; College of Bioscience and Biotechnology, Chubu University; Center for Mathematical Science and Artificial Intelligence, Chubu University

## Abstract

Complex chromosomal rearrangements (CCRs) are often observed in clinical samples from patients with cancer and congenital diseases but are difficult to induce experimentally. For generating animal models, these CCRs must be induced as desired, otherwise they cause profound genome instability and/or result in cell death. Here, we report the first success in establishing animal models for CCRs. The disruption of *Recql5*, which degrades RAD51 during DNA repair, successfully induces CRISPR/Cas9-mediated CCRs, establishing a mouse model containing triple fusion genes and megabase-sized inversions. Notably, some of these structural features of individual chromosomal rearrangements use template switching and microhomology-mediated break-induced replication mechanisms and are reminiscent of the newly described phenomenon “chromoanasynthesis.” Whole-genome sequencing analysis revealed that the structural variants in these mice caused only target-specific rearrangements. Thus, these data show that Recql5-deficient mice would be a novel powerful tool for analyzing the pathogenesis of CCRs, particularly chromoanasynthesis, whose underlying mechanisms are poorly understood.

## Introduction

Recent advances in bioinformatics technologies have led to the detection of complex chromosome rearrangements (CCRs) consisting of ≥3 chromosomal breaks in patients with cancer and congenital diseases^1^. These rearrangements caused by catastrophic cellular events can affect phenotype, thereby inducing a disease-promoting environment^2^. In particular, chromoanasynthesis, a recently discovered form of CCRs, is caused by erroneous DNA replication of a single chromosome through fork stalling and template switching (FoSTeS) and microhomology-mediated break-induced replication (MMBIR), which generate regions with complex rearrangements^3^. However, the pathogenic mechanisms underlying these diseases remain unclarified, and the establishment of appropriate animal models is essential for their elucidation. Although recent technologies, such as clustered regularly interspaced short palindromic repeats (CRISPR)-dependent base editing, prime editing, and DNA integration, have allowed for high-precision genome interrogation^4^, they have not yet been adapted to model CCRs in the germ line.

Here, we hypothesized that the efficient induction of CCRs could be achieved by manipulating the DNA repair pathway as accumulating evidence indicates that changes in DNA repair timing often accompany genomic rearrangements^2^. However, given the importance of DNA repair genes in genome maintenance, these strategies may have adverse consequences. Indeed, most genes involved in the DNA repair pathway are essential, and their homozygous disruption leads to embryonic lethality in mice^5^. However, *Recql5-*deficient mice were reported to live to adulthood^6^. RecQ protein-like 5 (RECQL5) helicases can displace the DNA repair protein RAD51 from single-stranded (ss)DNA and disassemble nucleoprotein filaments, thereby suppressing homology-directed repair (HDR)^6^. Transient accumulation of the homologous recombination and RAD51 has been reported in *Recql5*-deficient cells, which could alter DNA repair pathways, thereby contributing to chromosomal rearrangements^6^. Hence, we investigated whether CCRs might be induced in *Recql5*-deficient mice and succeeded in establishing CCRs model mice. Notably, a novel DNA repair system, FoSTeS/MMBIR, was involved in a CCR model mouse line.

## Results and Discussion

### *Recql5* mutant enables CRISPR/Cas9-mediated CCRs in mouse zygotes

Using a previously developed *in vivo* electroporation technique, called improved genome editing via oviductal nucleic acid delivery (*i*-GONAD)^7^ (Figure S1), we first established a mouse strain with a deletion of the RAD51-binding domain in RECQL5 (*Recql5*^*em1Cu*^) (Figure S2A, S2B). These *Recql5*^*em1Cu/em1Cu*^ mice were fertile and unexpectedly showed no overt signs of other diseases, such as tumorigenesis or inflammation (Figure S2C, S2D); five male and four female mice were observed for more than 60 weeks.

Second, we sought to investigate whether the *Recql5* mutant approach could be effectively utilized to generate inversion rearrangement mouse models. We recently succeeded in inducing a 7.67-Mb inversion in wild-type (WT) mice^8^ (Figure 1A). We induced this large inversion in *Recql5*^*em1Cu/ em1Cu*^ mice as well, with higher genome editing efficiency than that in WT mice (Figure 1B, 1I). Homozygous inversion mice, *In(15)*^*#6*^, were generated by breeding heterozygous males and females (Figure 1C), and exhibited a white-spotted phenotype due to disrupted *Adamts20* expression.

**Fig. 1.**
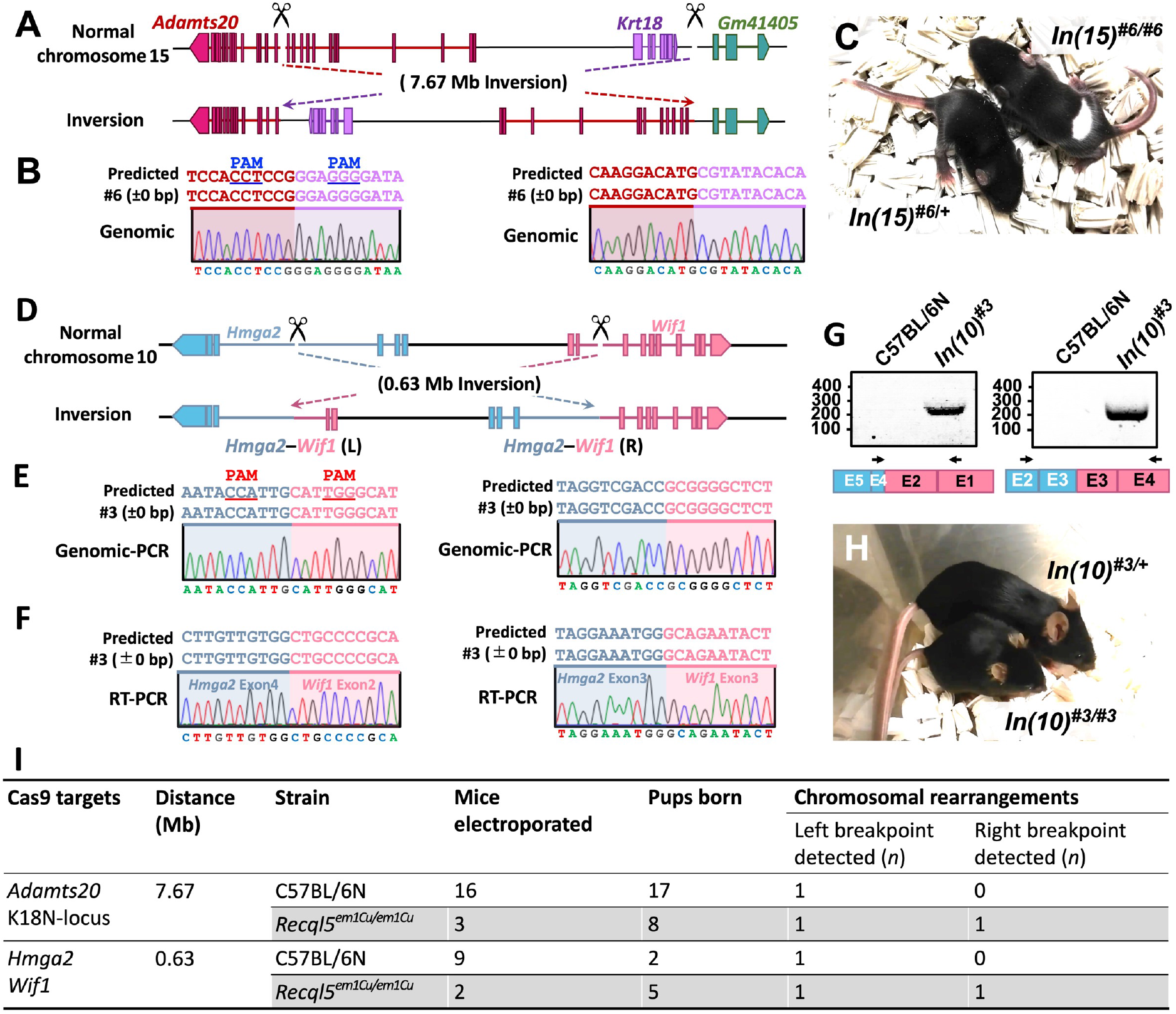
Chromosomal engineering of inversions using the *Recql5* mutant. (A) Schematic illustration of the chromosomal rearrangements created by an inversion between *Adamts20* and the K18N-locus in chromosome 15. (B) Alignment of sequences corresponding to the *Adamts20* and the K18N-locus genomic breakpoint junctions. (C) Generated inversion *In(15)*^*#6*^ on chromosome 15 shows a recessive white-spotted phenotype. (D) Schematic illustration of the chromosomal rearrangements created by an inversion between the *Hmga2* and *Wif1* genes in chromosome 10. (E) Alignment of sequences corresponding to the *Hmga2* and *Wif1* genomic breakpoint junctions. (F) Sanger sequences corresponding to the *Hmga2–Wif1* cDNA. (G) PCR amplification of the predicted fusion transcript. (H) Generated inversion, *In(10)*^*#2*^, on chromosome 10 showed a recessive pygmy phenotype. (I) Summary of the experimental efficiency of chromosomal inversions.

The human *HMGA2–WIF1* fusion gene, which was generated by the inversion on chromosome 12, was shown to activate the Wnt/β-catenin pathway and has been found in salivary gland tumors and breast adenomyoepitheliomas^9^. Likewise, we efficiently modeled the *HMGA2–WIF1* inversion on chromosome 10 in *Recql5*^*em1Cu/ em1Cu*^ mice (Figure 1D, 1E, and 1I), and the resulting mice, named *In(10)*^*#2*^, invariably harbored the *Hmga2–Wif1* inversion. These mice expressed the *Hmga2–Wif1* fusion gene (Figure 1F, 1G), and displayed a recessive pygmy phenotype due to the *Hmga2* mutation^10^ (Figure 1H). Taken together, these results indicate that the *Recql5* mutant approach can efficiently generate inversion mouse models.

Next, we attempted to apply this technique to produce CCR model mice. We selected an approximately 1.1-Mb region of mouse chromosome 10 containing *Hmga2, Wif1*, and *Rassf3* (Figure 2A), as these three genes are involved in human cancer and are mapped in a similar configuration in human chromosome 12. *RASSF3* is an important gene in p53-dependent apoptosis and functions as a tumor suppressor^11^. We designed gRNAs targeting these three genes and single-stranded oligodeoxynucleotides (ssODNs) that joined the chromosomal breakpoints, each of which had a sequence homologous to each junction point, such that two inversions were induced by the HDR process between the targeted regions and the homologous ssODNs. The 5′ and 3′ ends of the ssODNs were protected with two consecutive phosphorothioate-modified bases to improve the efficiency of HDR^12^. We then injected CRISPR/Cas9 ribonucleoproteins (RNPs) targeting these genes into pregnant females to generate chromosomal rearrangements^7^ (Figure S1). The generation of predicted chromosomal rearrangements was first confirmed by PCR of genomic DNA and then validated by sequencing the corresponding fusion transcript. Here, founder (F0) mice, in which a central breakpoint was detected, were defined as having induced CCRs and were used for subsequent analyses.

**Fig. 2.**
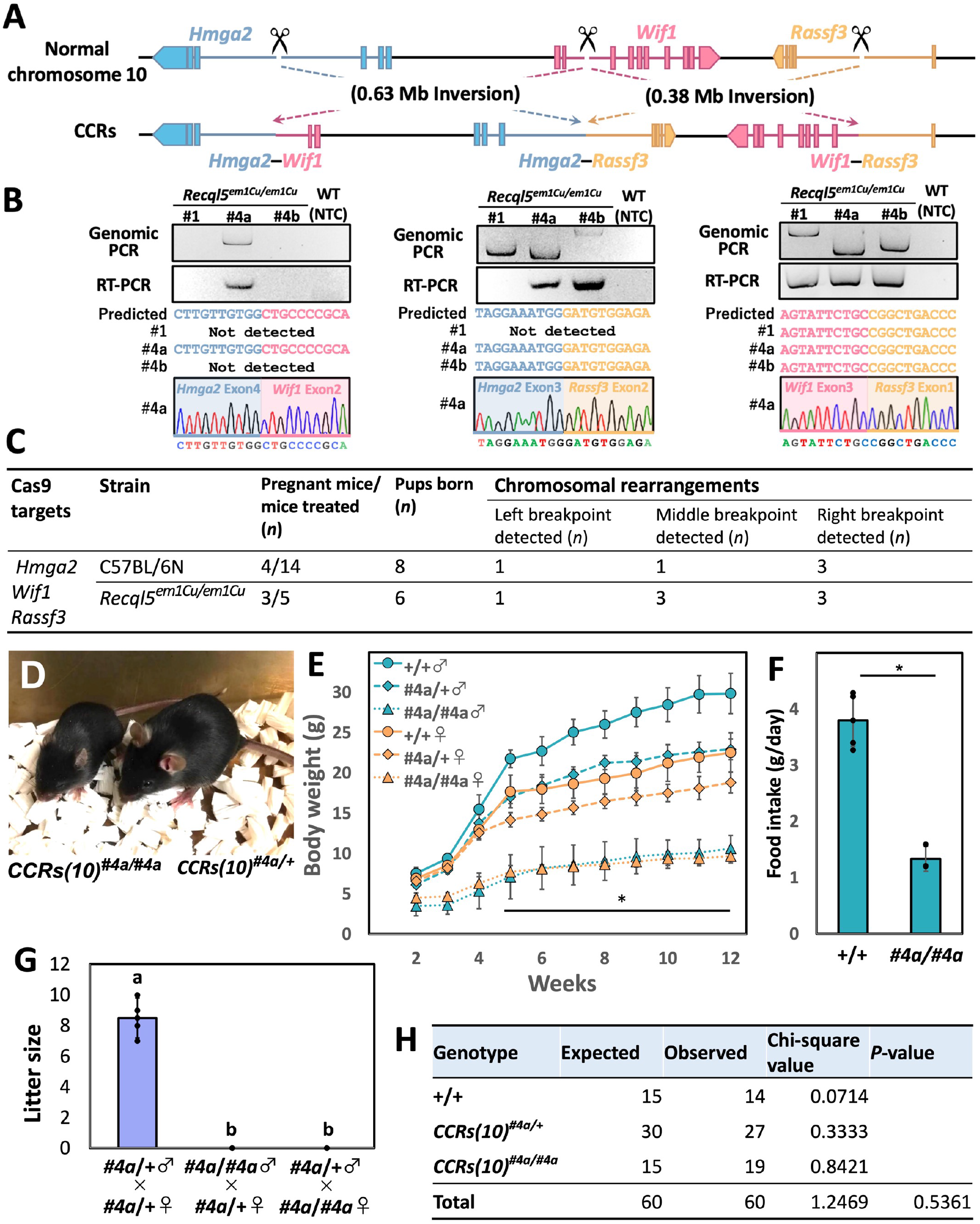
Induction of *Hmga2*–*Wif1, Hmga2*–*Rassf3*, and *Wif1*–*Rassf3* complex rearrangements in mouse zygotes using the *Recql5* mutant. (A) Schematic representation of the CCRs created between *Hmga2, Wif1*, and *Rassf3* in chromosome 10. (B) PCR amplification of the predicted breakpoint junction and fusion transcript. Sanger sequences corresponding to *Hmga2–Wif1, Hmga2–Rassf3*, and *Wif1–Rassf3* cDNA. NTC: Negative control. (C) Summary of the experimental efficiency of complex chromosomal rearrangements. (D) Appearance of heterozygous and homozygous *CCRs(10)*^*#4a*^ male mice at 6□weeks of age. (E) Growth curves of WT (+/+) (n = 11/10), heterozygous *CCRs(10)*^*#4a*^ (#4a/+) (n = 7/8), and homozygous *CCRs(10)*^*#4a*^ (#4a/#4a) (n = 9/12) mice (female/male). Error bars, mean□±□S.D. \**P*□<□0.05 (two-tailed Student’s *t*-test). (F) Food intake (g/day) of WT (+/+) and *CCRs(10)*^*#4a/#4a*^ male mice (20□weeks of age). Error bars, mean□±□S.D. \**P*□<□0.05 (two-tailed Student’s *t*-test). (G) Comparison of litter sizes. Error bars, mean□±□S.D. (n = 6). Different letters above the bars indicate significant differences at *P* < 0.05 by Tukey’s HSD. (H) Mendelian ratios of new-born mice from *CCRs(10)*^*#4a*^ heterozygous crossings.

In the *Recql5*^*em1Cu/ em1Cu*^ strain, we obtained six F0 pups via cesarean section and found that four had chromosomal rearrangements in the target locus, yielding three viable F0 CCR mice (Figure 2B, 2C: #1, #4a, #4b). By contrast, control WT (C57BL/6N) strains showed partial chromosomal rearrangements in three of the eight pups; however, we could not obtain the surviving founder, F0 (Figure 2C). Imprecise repair of double-strand breaks (DSBs) has the potential to be highly deleterious, owing to genomic instability, including the formation of chromosomal rearrangements^13^. Despite this seemingly difficult chromosomal rearrangement pattern, we confirmed the expression of the corresponding fusion transcript using reverse transcription (RT)-PCR and direct sequencing (Figure 2B). Homozygous *CCRs(10)*^*#4a*^ mice, in which all three fusion genes resulted in a recessive pygmy phenotype due to the *Hmga2* mutation^10^ (Figure 2D), including severe growth retardation and infertility (Figure 2E–2G), did not develop tumors within the timeframe of analysis. These findings suggest that the expression of the Hmga2–Wif1 fusion protein does not necessarily play a tumor promoting role in mice, although observations over a prolonged period are required to assess this possibility. The mating of heterozygous *CCRs(10)*^*#4a*^ mice resulted in homozygous, heterozygous, and WT mice born with the expected Mendelian inheritance (Figure 2H).

### Evaluation of genome-wide target specificity in the *Recql5* mutant

We characterized the genomic structure of the mouse strains in detail using whole-genome sequencing (WGS). The *CCRs(10)*^*#4a*^ strain showed multiple breakpoint junctions and was classified as deletion (red), duplication (green), and inversion (teal and blue) based on paired-ends with read depth changes (Figure 3A, 3B). Sequencing of the breakpoint revealed microhomology patterns and sister chromatid-containing templates, in which the added insertion was dependent on the Cas9 target site (Figure 3C). Notably, microhomologies seemed to be used to switch the nearby template during DSB repair stalling and collapse—a process termed FoSTeS and MMBIR^14,15,16^. This structural feature of individual chromosomes is reminiscent of the newly described phenomenon chromoanasynthesis, which is observed in tumors, as well as in patients with congenital diseases^3^. In addition to FoSTeS/MMBIR and tandem inversions, a 601-kb deletion was identified in the C*CRs(10)*^*#1*^ strain (Figure S3A–S3C). Consistent with this result, the C*CRs(10)*^*#1*^ strain did not express the fusion transcript corresponding to the middle of *Hmga2–Rassf3*, and no homozygotes were found (Figure 2B, Figure S3E, S3F). The *CCRs(10)*^*#4b*^ genome contained a 1,016-kb duplication harboring *Wif1* and *Rassf3* (Figure S4A–S4C). No discernible differences in phenotype were observed between the *CCRs(10)*^*#4b*^ and WT strains (Figure S4E, S4F).

**Fig. 3.**
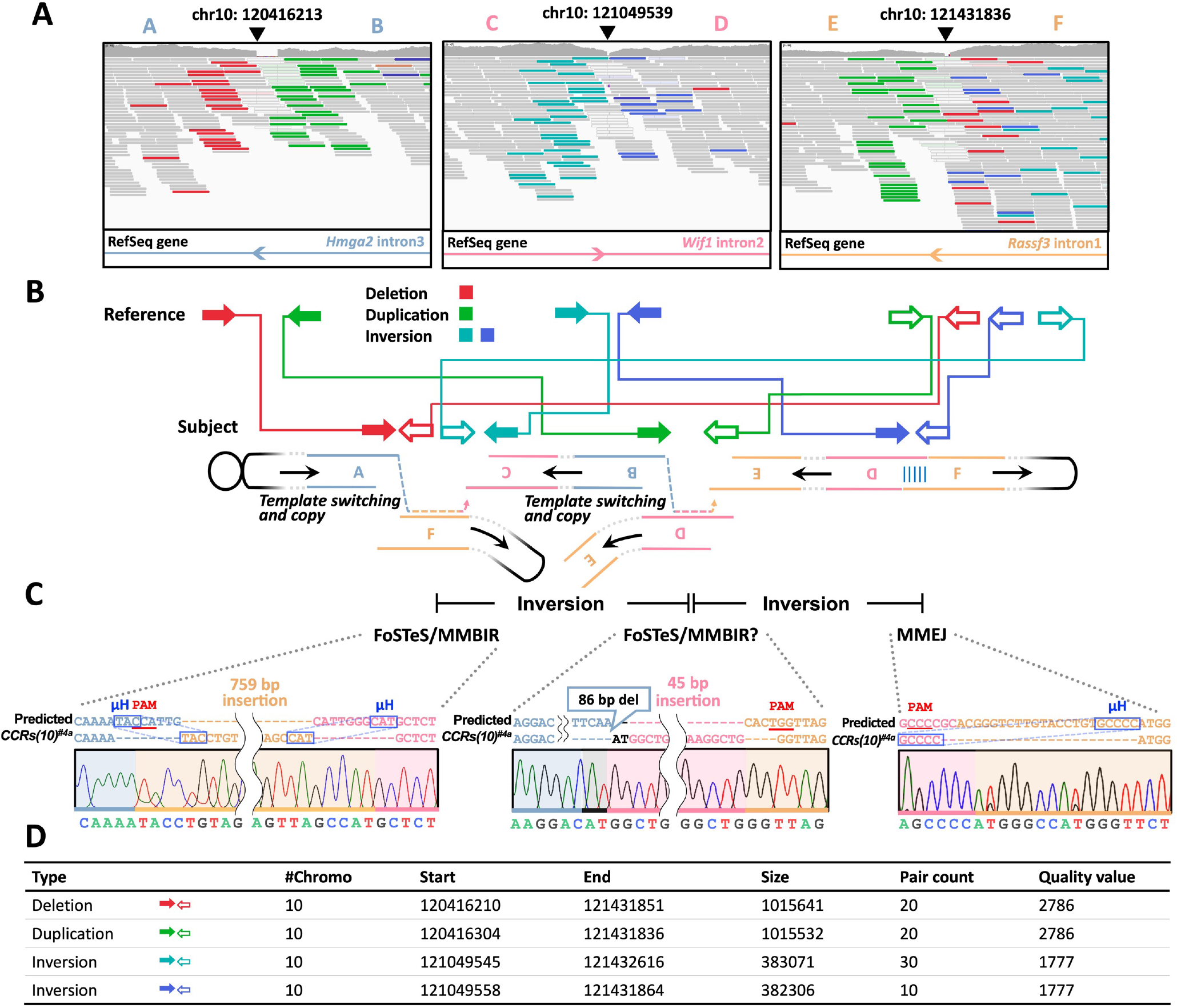
Validation of genome-wide target specificities in the *Recql5* mutant. (A) Whole-genome sequencing results. Interrogative genomic viewer (IGV) browser of CCRs data aligned to the mouse genome (mm10). gRNA cut sites are shown using black arrowheads. Deleted regions are shown in red; duplicated regions are shown in green; teal and blue reads indicate that they are mapped to the reverse strand. Normal reads are shown in gray. (B) Schematic indicating paired-end read interpretation in IGV for complex rearrangements. Read pairs are colored as per their interpretation in IGV. (C) Alignment of sequences from PCR products corresponding to the *Hmga2–Wif1, Hmga2–Rassf3*, and *Wif1–Rassf3* genomic breakpoint junctions. PAM, protospacer-adjacent motif; μH, microhomology. (D) Manta calls supporting chromosomal breakpoints.

Contrastingly, HDR appeared to repair two of the five breakpoints examined in the WT background, while the others were likely repaired by microhomology-mediated end joining or non-homologous end joining; however, no FoSTeS/MMBIR was observed (Table S1). A complex genome architecture confuses the DNA repair machinery and induces template-switching events driven by FoSTeS and MMBIR^3,17^. Thus, if the RAD51-filament is retained, loss of *Recql5* function may promote DNA repair machinery confusion and dictate the choice between the FoSTeS/MMBIR pathways.

These candidate rearrangement breakpoints were identified from WGS data using the Manta structural variation detection algorithm; however, no target-independent complex rearrangements were identified by comparing the test genomes^18^ (Figure 3D, Figure S3D, S4D). Thus, these results establish the efficiency and specificity of chromosomal engineering using the proposed approach.

### *Recql5* mutant mediates a broad pattern of chromosomal rearrangements

To test the broad applicability of the *Recql5* mutant approach, we generated CCRs on chromosome 2, which included topologically associating domains at the HoxD loci. These loci are necessary for the development of the proximal part of the limb, including the future arm and forearm^19^. To study the effects of the structural variants, we engineered CCRs comprising sequential inversions between *Atf2* neighborhood (*Atf2N*) and *Hoxd1N* and between *Hoxd1N* and *Nfe2l2N* (Figure 4A). In the *Recql5*^*em1Cu/em1Cu*^ strains, we screened F0 mice using PCR amplification and DNA sequencing of three junction points and detected chromosomal rearrangements in two of the four F0 mice (Figure 4B, 4C). In one of the possible CCR mouse lines, named F0-#2, two HDR-repaired junction points were detected, whereas the other junction point was repaired with structural changes that could not be amplified by PCR (Figure 4B). By contrast, the control C57BL/6N strain did not show any breakpoints in the five pups (Figure 4C). Although the RNP-based CRISPR/Cas system is characterized by high-efficiency germline transmission^20^, F0-#2 did not transmit the targeted CCRs to their 34 offspring. In additional studies, we examined DNA extracted from the testes and semen of F0-#2 animals; however, the semen samples did not exhibit a PCR signal between *Atf2N* and *Nfe2l2N* (Figure 4D). In our experience, F0-#2 was the first mouse line in which the mutation was not transmitted to the gametes using an RNP-based CRISPR/Cas system. Although we cannot accurately explain this transmission disturbance, asymmetric disjunction may result in a wide range of highly imbalanced gametes, many of which do not survive until the end of spermatogenesis^21^. Perhaps, breakpoints in intergenic regions may disturb the interactions between the promoter and transcriptional units with its *cis*-acting regulators through positional effects, thereby severely affecting gene expression^22^.

**Fig. 4.**
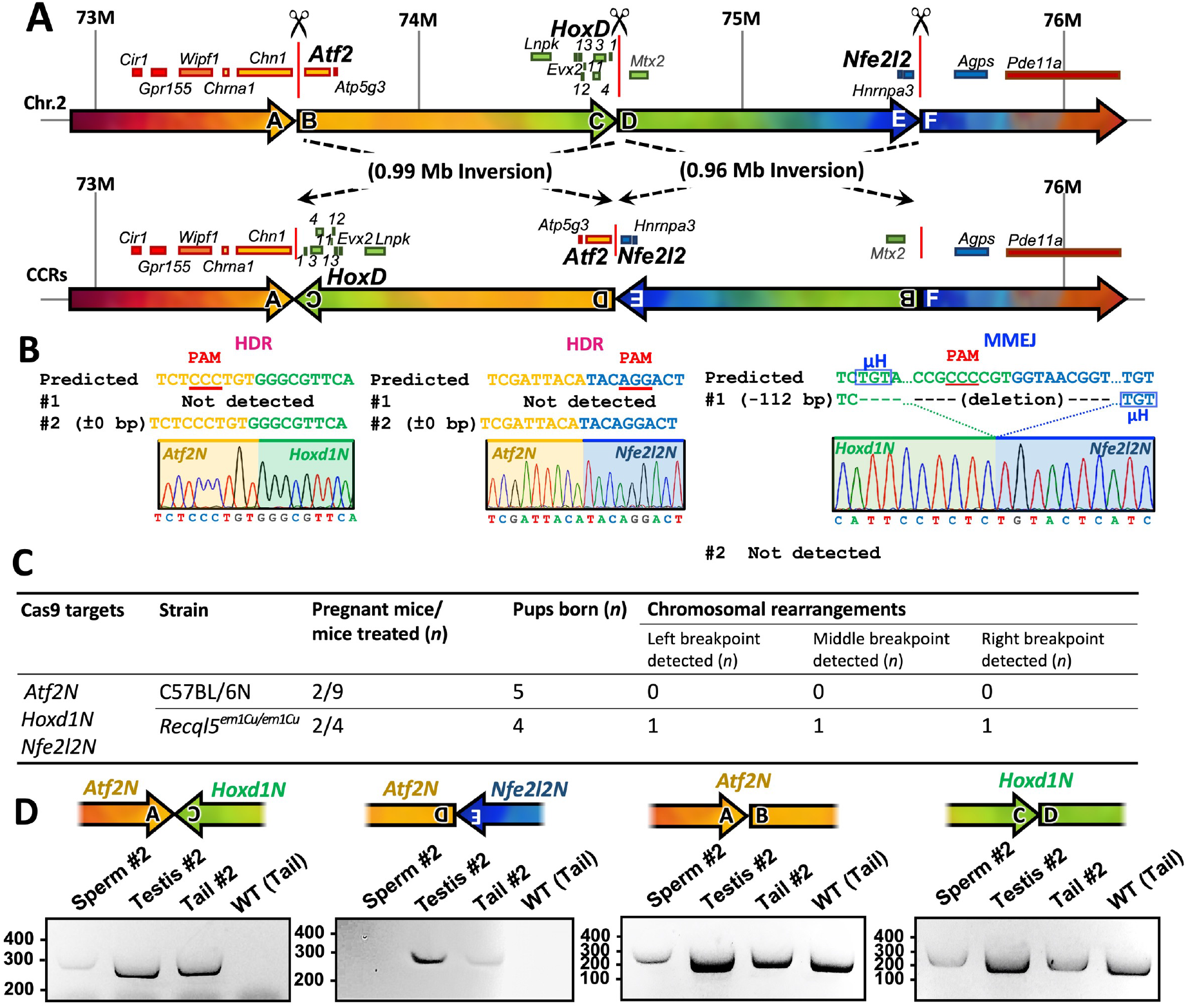
Complex chromosomal engineering in mouse zygotes at the HoxD loci using the *Recql5* mutant. (A) Schematic of the CCRs created between the *Atf2N, Hoxd1N*, and *Nfe2l2N* in chromosome 2. (B) Alignment of the sequences corresponding to the *Atf2N–Hoxd1N, Atf2N–Nfe2l2N*, and *Hoxd1N–Nfe2l2N* genomic breakpoint junctions. (C) Summary of the experimental efficiency of complex chromosomal rearrangements. (D) PCR amplification of the breakpoint junctions in CCR mouse sperm, testes, tail, and wild-type (WT).

To the best of our knowledge, this is the first study to successfully introduce CCRs into mouse zygotes using CRISPR-based methods. This approach opens new avenues for investigating the role of chromosomal rearrangements in diseases and provides a powerful tool for engineering genetic modifications in various organisms.

### Limitations of the study

Here, we describe a new genome editing method using a *Recql5* mutant mouse and show that it can efficiently induce various types of chromosomal rearrangements, including CCRs and inversions, in mouse zygotes. Although this technique has considerable potential, it is also associated with unexpected DSB repair mechanisms such as FoSTeS/MMBIR. Thus, the structural features of induced rearrangements are likely to depend on a variety of factors, such as features specific to the targeted genomic regions, the number of breakpoints, and the unique properties of mouse lines.

In FoSTeS and MMBIR models, the absence of a homologous template during HDR possibly results in the activation of microhomology pairing repair of broken ends, a more error-prone DNA repair. Thus, these factors need to be carefully considered during the application of such techniques to therapeutics.

## Author contributions

S.I. designed and performed the experiments and drafted the manuscript. M.N. performed the animal experiments. T.I. supervised the studies and revised the manuscript. All authors have read and approved the final manuscript.

## Acknowledgments

The authors thank the laboratory members for their support with animal care and experiments. This study was supported by JSPS KAKENHI for a Grant-in-Aid for Scientific Research (B) (to S.I., Grant Number 21H02395), The Science Research Promotion Fund 2022 (to S.I., Grant Number 29), and The Research Institute of Life and Health Sciences, CHUBU University Grant 2021 (to S.I.).

## Methods

### Experimental model and study participant details

C57BL/6NCrSlc mice (Japan SLC, Shizuoka, Japan) were used in this study. The animals were housed at a constant temperature (22□±□2 °C) and humidity (50□±□10%), with a 12-h light/12-h dark cycle. All animal experiments were approved by the Institutional Animal Care and Use Committee of Chubu University (Permit Number #202110033) and were conducted in accordance with the institutional guidelines.

### CRISPR RNP and ssODN preparation

The CRISPR guide RNAs were designed using CHOPCHOP^23^ (http://chopchop.cbu.uib.no/; Table S2). CRISPR RNP consists of Alt-R S.p. Cas9 Nuclease 3NLS (Integrated DNA Technologies, Coralville, IA, USA) and a custom guide RNA (crRNA): universal structural RNA (tracrRNA) duplex (Integrated DNA Technologies). crRNA and tracrRNA were heated to 95 °C for 10□min and slowly cooled to 25 °C. This crRNA:tracrRNA duplex and the Alt-R S.p. Cas9 Nuclease 3NLS were incubated at 25 °C for 10□min to form the RNP complex. The ssODNs were manufactured by Eurofins Genomics (Tokyo, Japan) and were designed to join two DNA sequences so that the junction would be positioned at the center of the predicted cleavage sites, which were located within 3□bp of the PAM sequences (Table S3). The 5′ and 3′ ends of the ssODNs were protected with two consecutive phosphorothioate-modified bases (*) to improve the HDR efficiency^12^ (Table S3).

### *i*-GONAD method

C57BL/6NCrSlc female mice (8–12-week-old) in estrus were mated with males (8–24-week-old) at 15:00–18:00 h. The presence of copulation plugs was confirmed the next morning via visual inspection, and plug-positive mice were subjected to *i*-GONAD experiments, as previously described^7^. The following CRISPR solutions were used: 540□ng/μL Alt-R S.p. Cas9 Nuclease 3NLS, 33□μM upstream and downstream crRNA:tracrRNA, and 0.05% Fast Green FCF (Wako, Osaka, Japan) marker diluted in Opti-MEM (Thermo Fisher Scientific, Waltham, MA, USA). Prior to electroporation, females were anesthetized with a mixture of medetomidine (0.75□mg/kg), midazolam (4□mg/kg), and butorphanol (5□mg/kg). The CRISPR mixture (1□μL) was injected into the oviductal lumen upstream of the ampulla using a glass micropipette including a vertical capillary puller (NARISHIGE, Tokyo, Japan). The CRISPR mixture was injected using a FemtoJet 4i microinjector (Eppendorf, Hamburg, Germany) with the following settings: pi: 100 hPa, ti: 0.2 s, and pc: 0 hPa. After injection of CRISPR solutions, the oviduct regions were clamped using tweezer electrodes (LF650P3; BEX, Tokyo, Japan), and electroporation was performed as previously described^7^ using a CUY21EDIT II (BEX, Tokyo, Japan). The following parameters were used for electroporation: square (mA), (+/−), Pd V: 60 V or 80 V, Pd A: 200 mA, Pd on: 5.00 ms, Pd off: 50 ms, Pd N: 3, Decay: 10%, DecayType: Log. Thereafter, we placed the oviducts back in their original location and sutured the incisions. Post operation, atipamezole hydrochloride (0.75□mg/kg) was intraperitoneally injected to reverse the effects of medetomidine.

### Analysis of CRISPR/Cas9-engineered mice

To screen for CRISPR/Cas9-induced mutations, genomic DNA was isolated from the tails or ears of founder mice using lysis buffer [100□mM NaCl, 200□mM sucrose, 10□mM ethylenediaminetetraacetic acid, 300□mM Tris (pH 8.0), and 1% sodium dodecyl sulfate], and DNA was examined by PCR amplification. The obtained PCR products were purified using NucleoSpin Gel and PCR Cleanup kit (Takara Bio, Shiga, Japan) and sequenced directly or were cloned into the pTAC-1 vector (Biodynamics, Tokyo, Japan), and the sequences of individual clones were determined using Sanger sequencing (Eurofins Genomics). The PCR primers used for genotyping are listed in Table S4.

### RT-PCR

RT-PCR was performed using total RNA. Total RNA was isolated from ear tissue using ISOSPIN Cell & Tissue RNA (Nippon Gene, Tokyo, Japan). Template cDNA was obtained using ReverTra Ace qPCR RT Master Mix (Toyobo, Osaka, Japan). The RT-PCR products were treated with NucleoSpin Gel and PCR Cleanup kit (Takara Bio) and directly analyzed using Sanger sequencing (Eurofins Genomics). The primers used for RT-PCR are listed in Table S4.

### Growth analysis

From 0 to 12 weeks after birth, the weights (g) of *CCRs(10)*^*#4a/#4a*^, *CCRs(10)*^*#4a/+*^, and WT (+/+) mice were recorded for growth curve analysis.

### Food intake analysis

Mice were housed individually, and food intake was measured in terms of grams of diet consumed per day.

### Mating test

Upon sexual maturation, male mice were caged with two females for at least 8 weeks. During the mating test, pups were counted for litter size measurements and their tails or ears were biopsied for genotyping.

### Whole-genome sequencing analysis

Total genomic DNA was extracted using a DNeasy Blood & Tissue Kit (Qiagen, Hilden, Germany). The library was prepared using TruSeq Nano DNA (Illumina, CA, USA) according to the manufacturer’s library quantification protocol. Sequencing was performed as pair-end (150 bp) using an Illumina NovaSeq6000 (Illumina) by Macrogen sequencing service (Macrogen Inc., Seoul, Korea). Whole-genome sequencing analysis and genomic coordinates provided in this manuscript are based on the GRCm38/mm10 assembly. The sequencing data were then aligned in BAM format to the mouse reference genome (mm10) using the Integrative Genomics Viewer (IGV-version 2.3.93)^24^. Structural rearrangements and breakpoints were identified using Manta Structural Variant Caller^18^ (version 1.3.1). Breakpoints were visually inspected using IGV (IGV-version 2.3.93) to confirm the presence of split and spanning reads. The whole-genome sequencing data reported in this paper have been deposited into the DNA Data Bank of Japan (DDBJ) Sequence Read Archive (DRA) (https://ddbj.nig.ac.jp/resource/bioproject/PRJDB15772).

### Quantification and statistical analysis

The Student’s *t*-test (two-tailed test) was used for body weight, food intake, and mating analyses. Data are presented as the mean ± standard deviation. For Mendelian genotype ratios of progeny obtained from sibling mating between CCRs mice, the chi-square test was performed using Excel version 16.36 (Microsoft, Redmond, WA, USA). Statistical comparisons were made using Tukey’s honestly significant difference test. The threshold for statistical significance was *P*□<□0.05.

