## Supplemental Information for "A *Recql5* mutant enables complex chromosomal engineering of mouse zygotes"

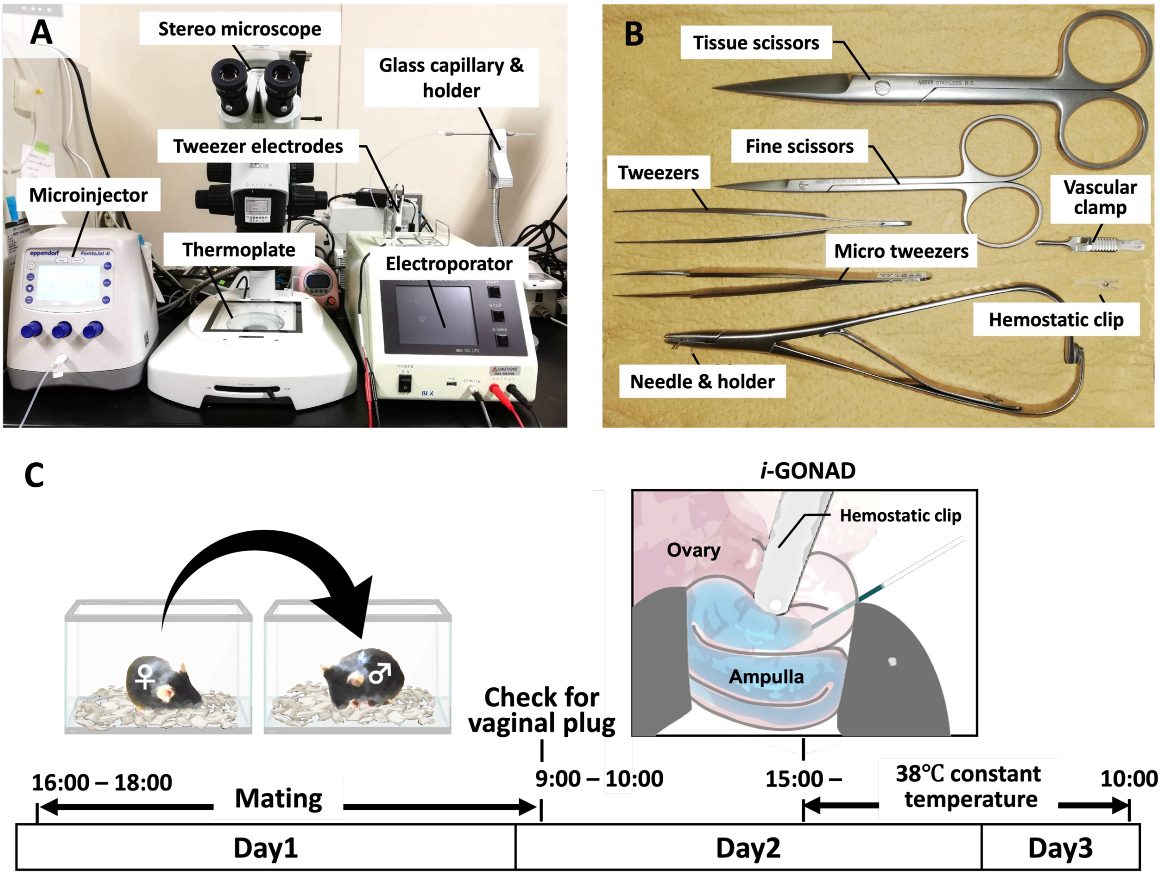


**Figure S1. Equipment for *i*-GONAD of mouse zygotes (related to STAR Methods).** (A) The instruments used for both techniques—injection and electroporation. (B) Surgical equipment. (C) Experimental procedures for induction of CCRs using *i*-GONAD.

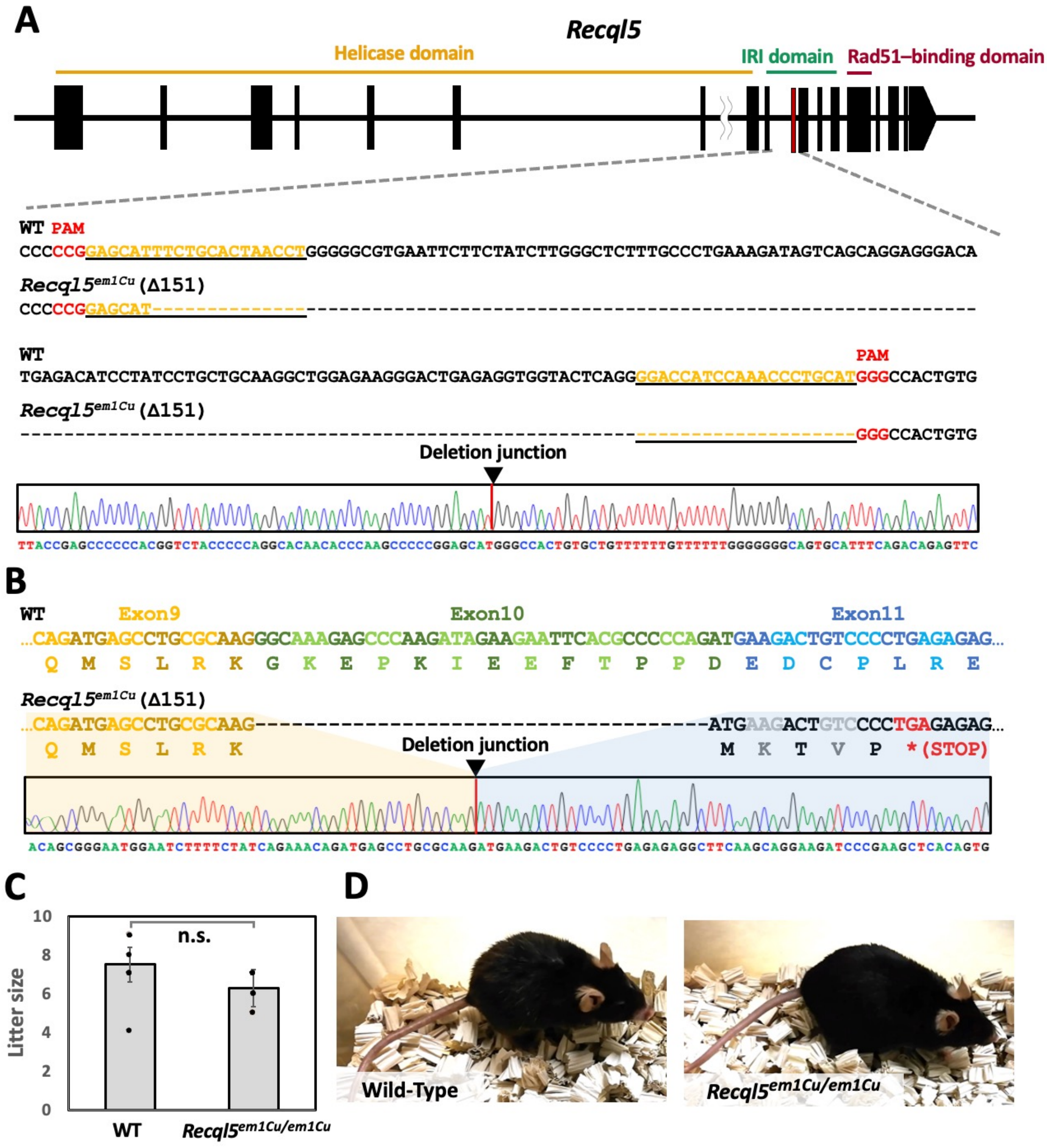


**Figure S2. Generation and phenotype of *Recql5* mutant mouse (related to Figure 1, 2, and 4).**

(A) Schematic representation of *Recql5* gene deletion. The gRNA sequence is underlined in black. The PAM sequence is indicated in red. (B) Alignment of sequences corresponding to the *Recql5* cDNA. The *Recql5*$^{em1Cu}$ caused a frameshift mutation. (C) Comparison of the litter sizes of wild-type (WT) (n = 7) and *Recql5*$^{em1Cu/em1Cu}$ mice (n = 18). n.s., not significant. (D) Images of representative post-natal year 2 wild-type (WT) (left) and *Recql5*$^{em1Cu/em1Cu}$ (right) male mice.

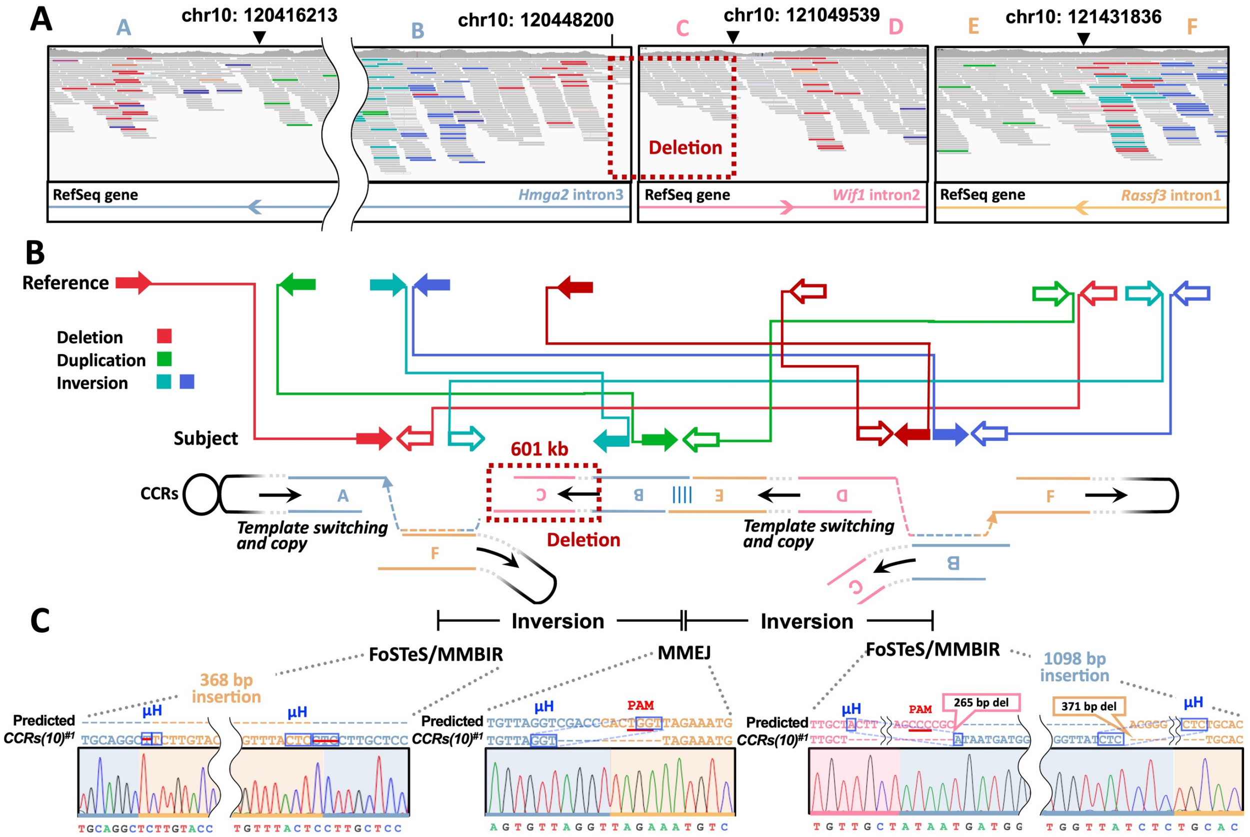


**D**

| Type | | #Chromo | Start | End | Size | Pair count | Quality value |
|---|---|---|---|---|---|---|---|
| Deletion | →⇦ | 10 | 120415542 | 121431846 | 1016304 | 11 | 717 |
| Deletion | →⇦ | 10 | 120448107 | 121049828 | 601721 | 11 | 77 |
| Duplication | →⇦ | 10 | 120416224 | 121431833 | 1015609 | 4 | 717 |
| Inversion | →⇦ | 10 | 120447014 | 121432214 | 985200 | 11 | 1167 |
| Inversion | →⇦ | 10 | 120447015 | 121432215 | 985200 | 21 | 1167 |

**E**

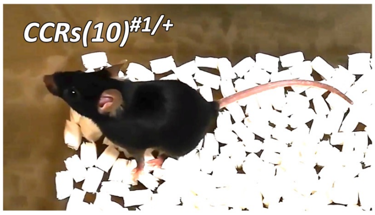


**F**

| Genotype | Expected | Observed | Chi-square value | *P*-value |
|---|---|---|---|---|
| +/+ | 8.5 | 8 | 0.02941 | |
| *CCRs(10)*$^{\#1/+}$ | 17 | 26 | 4.7647 | |
| *CCRs(10)*$^{\#1/\#1}$ | 8.5 | 0 | 8.5 | |
| Total | 34 | 34 | 13.2941 | 0.001297 |

**Figure S3. Validation of genome-wide target specificities in *CCRs(10)*$^{\#1}$ mice (related to Figure 3).**

(A) Whole-genome sequencing results. IGV browser image of *CCRs(10)*$^{\#1}$ data aligned to the mouse genome (mm10). gRNA cut sites are shown by black arrowheads; deleted regions are shown in red; duplicated regions are shown in green; teal and blue reads indicate that they are mapped to the reverse strand; normal reads are shown in gray. (B) Schematic illustration indicating paired-end read interpretation in IGV for complex rearrangements. Read pairs are colored as per their interpretation in IGV. (C) Alignment of sequences from PCR products corresponding to the *Hmga2–Wif1*, *Hmga2–Rassf3*, and *Wif1–Rassf3* genomic breakpoint junctions. (D) Manta calls supporting chromosomal breakpoints. (E) Image of representative heterozygous *CCRs(10)*$^{\#1}$ mouse. (F) Mendelian ratios of new-born mice from *CCRs(10)*$^{\#1}$ heterozygous crossings.

A

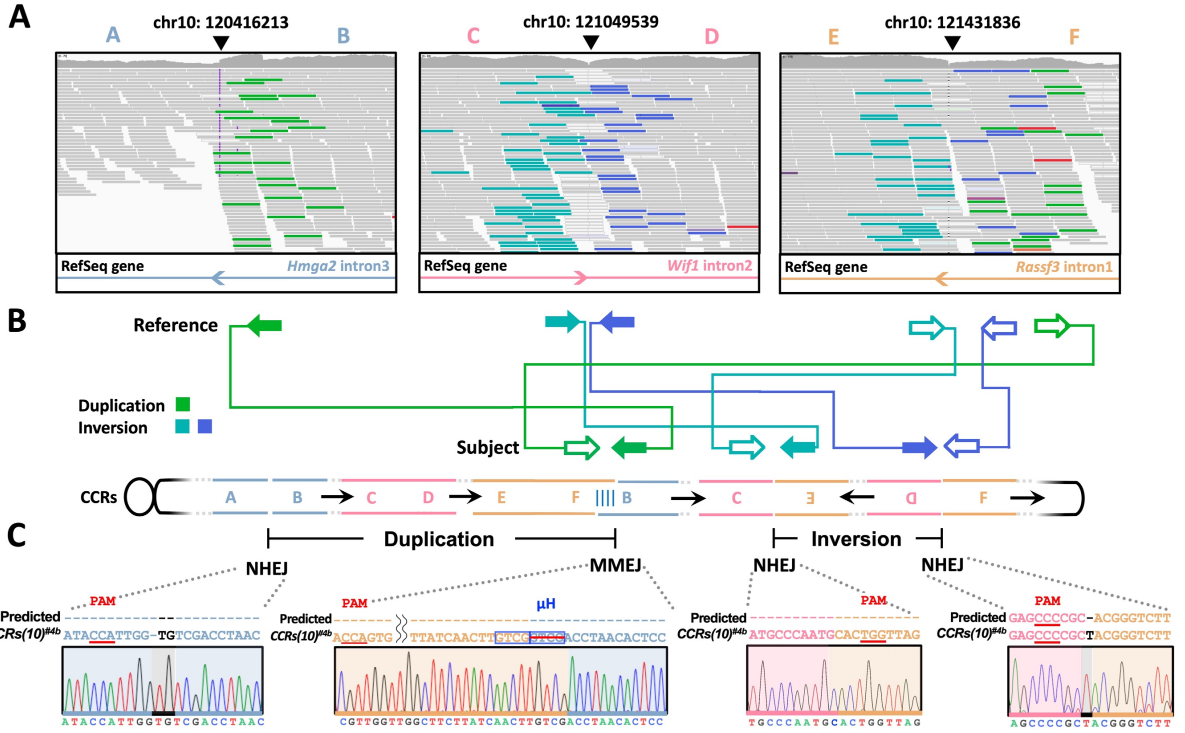


D

| Type | | #Chromo | Start | End | Size | Pair count | Quality value |
|---|---|---|---|---|---|---|---|
| Duplication | ➡⇦ | 10 | 120416220 | 121432457 | 1016237 | 30 | 959 |
| Inversion | ➡⇦ | 10 | 121049493 | 121431816 | 382323 | 13 | 2635 |
| Inversion | ➡⇦ | 10 | 121049555 | 121431840 | 382285 | 58 | 1777 |

E

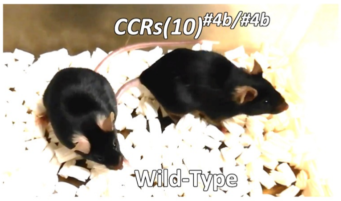


F

| Genotype | Expected | Observed | Chi-square value | *P*-value |
|---|---|---|---|---|
| +/+ | 5 | 6 | 0.2 | |
| *CCRs(10)*$^{\#4b/+}$ | 10 | 7 | 0.9 | |
| *CCRs(10)*$^{\#4b/\#4b}$ | 5 | 7 | 0.8 | |
| Total | 20 | 20 | 1.9 | 0.38674102 |

**Figure S4. Validation of genome-wide target specificities in *CCRs(10)*$^{\#4b}$ mice (related to Figure 3).**

(A) Whole-genome sequencing results. IGV browser image of *CCRs(10)*$^{\#4b}$ data aligned to the mouse genome (mm10). gRNA cut sites are shown by black arrowheads; deleted regions are shown in red; duplicated regions are shown in green; teal and blue reads indicate that they are mapped to the reverse strand; normal reads are shown in gray. (B) Schematic illustration indicating paired-end read interpretation in IGV for complex rearrangements. Read pairs are colored as per their interpretation in IGV. (C) Alignment of sequences from PCR products corresponding to the *Hmga2–Wif1*, *Hmga2–Rassf3*, and *Wif1–Rassf3* genomic breakpoint junctions. (D) Manta calls supporting chromosomal breakpoints. (E) Images of representative wild-type (WT) and homozygous *CCRs(10)*$^{\#4b}$ mice. (F) Mendelian ratios of new-born mice from *CCRs(10)*$^{\#4b}$ heterozygous crossings.

**Table S1. Sequences of the CCR genomic breakpoints obtained in wild-type mice (related to Figure 3).**

| Founder # | Left breakpoint | Middle breakpoint | Right breakpoint |
|---|---|---|---|
| #2 | Not detected | µH PAM<br>Predicted GTGTTAGGTCGACCCACTGGTTAGAAATGTCG<br>#2 GTGTTAGGT------------TAGAAATGTCG | PAM<br>Predicted AGAGCCCCGCACGGGTCTTG<br>#2 AGAGCCCCGCACGGGTCTTG |
| #5 | Not detected | Not detected | PAM<br>Predicted AGAGCCCCGCACGGGTCTTG<br>#5 AGAGCCCCGCACGGGTCTTG |
| #9 | PAM<br>Predicted AAATACCATTGCATTGGGCATGCTCTATTGAAGGCTGAGTCCAGAGCTCTCCATTGGGC<br>#9 AAATACCATT--------------------------------------------TGGGC | Not detected | PAM<br>Predicted GAGCCCCGC-----ACGGGTCTTGTA<br>#9 GAGCCCCG-ATTTT----GTCTTGTA |

**Table S2. gRNAs used in the present study (related to STAR Methods).**

| Target loci | Genomic location (mm10/GRCm38) | Target Sequences (PAM) | CHOPCHOP (https://chopchop.cbu.uib.no/) | | | | |
|---|---|---|---|---|---|---|---|
| | | | Number of mismatches | | | | |
| | | | 0 | 1 | 2 | 3 | Efficiency |
| Recql5 | chr11:115896078 | AGGTTAGTGCAGAAATGCTC (CGG) | 0 | 0 | 2 | 6 | 46.87 |
| Recql5 | chr11:115896218 | GGACCATCCAAACCCTGCAT (GGG) | 0 | 0 | 0 | 2 | 57.44 |
| Adamts20 | chr15:94347708 | TCGTGTTCAAGGACATGCGG (AGG) | 0 | 0 | 1 | 3 | 71.44 |
| K18N-locus | chr15:102034830 | GAAGCCCTGTGTATACGGGA (GGG) | 0 | 0 | 1 | 4 | 61.49 |
| Hmga2 | chr10:120416213 | GGAGTGTTAGGTCGACCCAA (TGG) | 0 | 0 | 0 | 0 | 67.29 |
| Wif1 | chr10:121049539 | ATAGAGCATGCCCAATGGCG (GGG) | 0 | 0 | 0 | 0 | 60.84 |
| Rassf3 | chr10:121431836 | ACAGGTACAAGACCCGTCAC (TGG) | 0 | 0 | 0 | 2 | 59.01 |
| Atf2N | chr2:73802063 | AGCAAGGGTCGATTACAACA (GGG) | 0 | 0 | 0 | 0 | 70.21 |
| HoxD1N | chr2:74789654 | TAAAGCACTGAACGCCCACG (GGG) | 0 | 0 | 0 | 0 | 72.43 |
| Nfe2l2N | chr2:75750986 | TCAGGGTGACCGTTACCTAC (AGG) | 0 | 0 | 0 | 0 | 57.25 |

### Table S3. ssODNs used in the present study (related to STAR Methods).

| Target loci | Target Sequences (5′–3′) |
|---|---|
| Adamts20–K18N-locus (Left) | CGTACTCGGGCCGGTCACAGAGCCGGGCTGTGCTCTTGATTCCACCTCCGGGAGGGGATAATCAGTCAGGTGCCTCGGCTCAGGTTTCTCGGGTCAGCTT |
| Adamts20–K18N-locus (Right) | GATGGCGAGTGGGGACCATGGGGACCCTACAGCTCGTGTTCAAGGACATGCGTATACACAGGGCTTCGCAGTTCCCAGGGCTCTTACGCATTTGATCCTC |
| Hmga2::Wif1 (Left) | CGTTGCAGCATGTAGGTGTGGTAGCATCTGCACCCACCAAAATACCATTGCATTGGGCATGCTCTATTGAAGGCTGAGTCCAGAGCTCTCCATTGGGCAT |
| Hmga2::Wif1 (Right) | GAAGCACGAAGAAAAGAGTTTAAAAATTCAAAGGGAGTGTTAGGTCGACCGCGGGGCTCTCAGTAAGGCGTGGTTTTTTAAAAAAATCTTTTTTGGGTTA |
| Hmga2–Wif1 (Left) | C*G*TTGCAGCATGTAGGTGTGGTAGCATCTGCACCCACCAAAATACCATTGCATTGGGCATGCTCTATTGAAGGCTGAGTCCAGAGCTCTCCATTGGGC*A*T |
| Hmga2–Rassf3 (Middle) | G*A*AGCACGAAGAAAAGAGTTTAAAAATTCAAAGGGAGTGTTAGGTCGACCCACTGGTTAGAAATGTCGGGATTGGTTGCCCCCTGCCTCCTAGGCACT*G*G |
| Wif1–Rassf3 (Right) | T*A*ACCCAAAAAAGATTTTTTTAAAAAACCACGCCTTACTGAGAGCCCCGCACGGGTCTTGTACCTGTAGCCCCATGGGCCATGGGTTCTTGGCTAATT*T*T |
| Atf2N–Hoxd1N (Left) | C*A*GCTCAAACAACATAATGGCAGCAGGGAAGGACACCCTAGTCTCCCTGTGGGCGTTCAGTGCTTTAATTCCGTGTTTGTCACAGCGATTGCTTGACT*C*T |
| Atf2N–Nfe2l2N (Middle) | A*G*GCTACAAGACAAAAAGACCACAGGGAATTCTAGCAAGGGTCGATTACATACAGGACTGTGGGACTAGTGAGTGGGATTCGGGCGCAGAAGTAGCTA*T*G |
| Hoxd1N–Nfe2l2N (Right) | T*T*ATTATTCATTCCTCTCTGTACTTCACCTGATATTGAACTCCGCCCCGTGGTAACGGTCACCCTGACTTTTAATGTCATAGATTGATTTCATCAGAT*G*T |

**Table S4. Primers used in the present study (related to STAR Methods).**

| Target loci | Genomic location (mm10/GRCm38) | Sequences (5'–3') |
|---|---|---|
| Genotyping primers | | |
| Recql5-F | chr11: 115895976 to 115895997 | TCTTCATCTGAGAAAAGGGAGG |
| Recql5-R | chr11: 115896306 to 115896327 | CTGGAGTACAAGGCCAGAGACT |
| Recql5-Exon9-F | chr11: 115896851 to 115896872 | TGATGAAGGTTCTGGAGGTAGC |
| Recql5-Exon11-R | chr11: 115895792 to 115895813 | TGGTTACTGATCAGAGCCTCCT |
| Adamts20-F | chr15: 94347562-94347583 | TCTCTGAAACTCGCAGACTGAC |
| Adamts20-R | chr15: 94347823-94347844 | TTCCTGTGTGTGTGCTTCTTCT |
| K18N-locus-F | chr15: 102034680-102034700 | GGCATCAAATGTGTCTTCTCA |
| K18N-locus-R | chr15: 102034945-102034966 | TGATGTCTGTGGCCTTTACTGT |
| Hmga2-F | chr10: 120415727 to 120415748 | CCCCAGGGAAGGTAAATAATGT |
| Hmga2-R | chr10: 120416637 to 120416658 | TTGCTCTGGACAACATTCATTC |
| HMGA2-Exon4-F | chr10: 120374678 to 120374700 | TTCTTCTGAACGACTTGTTGTGG |
| HMGA2-Exon2-R | chr10: 120473262 to 120473285 | CAAAAACAAGAGCCCCTCTAAAGC |
| Wif1-F | chr10: 121049111 to 121049132 | TGACCCTCCACCATTAAAATTC |
| Wif1-R | chr10: 121049971 to 121049992 | AGATCATTGCTGGGAAGAAGAG |
| WIF1-Exon1-F | chr10: 121034441 to 121034462 | GGAGAGCTTGTACCTGTGGATC |
| WIF1-Exon4-R | chr10: 121082144 to 121082164 | CGTCTTGTTTGCCGAGACACG |
| Rassf3-F | chr10: 121431397 to 121431418 | GAAGACAGAGACAAACCATGCC |
| Rassf3-R | chr10: 121432300 to 121432321 | CTCTTTGTGGCACTTCTGTGTC |
| RASSF3-Exon2-F | chr10: 121417140 to 121417161 | TTGATTTCCTCTTTGCTGAGGT |
| RASSF3-Exon1-R | chr10: 121476034 to 121476054 | GACCTCCTTCTTCAGGAGAGC |
| Atf2N-F | chr2: 73802000 to 73802021 | CTCTTCGCAAGACACATTTCAG |
| Atf2N-R | chr2: 73802244 to 73802265 | CAGGCGTATTGGAAGACTTAGG |
| HoxD1N-F | chr2: 74789496 to 74789517 | GAGTGAACTGCAAGGAGGTTTC |
| HoxD1N-R | chr2: 74789720 to 74789741 | CCAGGAACTTCTCTGCTTCTCT |
| Nfe2l2N-F | chr2: 75750911 to 75750932 | TTCCAAATCAAGAGTGGAACAA |
| Nfe2l2N-R | chr2: 75751125 to 75751146 | AAACTAAACCACACCACAGCCT |
